# A multi-site MicroLED Optrode Array for Neural Interfacing

**DOI:** 10.1101/480582

**Authors:** Niall McAlinden, Yunzhou Cheng, Robert Scharf, Enyuan Xie, Erdan Gu, Martin D. Dawson, Christopher Reiche, Rohit Sharma, Prashant Tathireddy, Loren Rieth, Steve Blair, Keith Mathieson

## Abstract

We present an electrically addressable optrode array capable of delivering light to 181 sites in the brain, each providing sufficient light to optogenetically excite hundreds of neurons *in vivo*, developed with the aim to allow behavioural studies in large mammals. The device is a glass microneedle array directly integrated with a custom fabricated microLED device, which delivers light to 100 needle tips and 81 interstitial surface sites, giving 2-level optogenetic excitation of neurons *in vivo*. Light delivery and thermal properties are evaluated, with the device capable of peak irradiances > 80 mW/mm^2^ per needle at 50 ms pulse widths with tissue temperature increase less than 1 ^°^C. Future designs are explored through optical and thermal modelling and benchmarked against the current device.

## 1. Introduction

In a little over 10 years optogenetics has established itself as a key technique for modulating neuronal activity [1]. It has been used in ground breaking experiments allowing neuroscientists to de-code neuronal circuits [2, 3] and examine the behavioural consequences of activity in certain neuronal populations [4]. The rapid growth of the field has been driven by significant developments in both genetically encoded light-sensitive ion channels (such as channelrhodopsin, ChR2) and methods for transmitting light into neuronal tissue with the required intensity, spatial resolution and temporal characteristics.

More recently, optogenetic techniques have been applied in the non-human primate [5]. The non-human primate brain is the system closest to the human brain and plays an important role in our understanding of neural computation, cognition, and behaviour. Validation of optogenetic techniques in the non-human primate may allow for a better understanding of neural disorders which could lead to clinical applications. However, differences in scale between the rodent and primate brain mean that current devices used to deliver light for optogenetics may not be suited for primate studies.

The main difficulty in coupling light into the brain is scattering in brain tissue. This problem is exacerbated as most opsins require blue or blue-green light to function, which experience high tissue scattering. There are two solutions to this problem: an implantable device that delivers light directly to the region of interest, or a variety of methods allowing for excitation of opsins with infrared (IR) light where scattering is reduced [6-8]. Although using IR light is an attractive proposition, the lack of available opsins at this wavelength or the need for complex optical systems, if using 2-photon methods, limits applicability [6].

To deliver light deeper into brain tissue, in localised volumes, several different devices for optogenetics have been designed. The simplest and most common approach is using a fibre optic cannula [9], which allows for a single large area optogenetic excitation site. Light is normally coupled to the cannula using a fibre optic, where the resultant tethering can restrict animal movement, affecting behaviour. However, options are becoming available that couple an LED directly to the cannula and allow for more natural animal movement [10]. This type of device is suitable for non-human primate studies where only a single excitation source is required.

Another device used in optogenetics consists of a probe structure with integrated microLEDs [11-15]. These devices allow for 10’s – 100’s of excitation sites, with each exciting a small volume of tissue (<0.01 mm^3^). They are electrically addressed, meaning there is potential for making a scalable, wireless device. Integrated microLED probes of this type provide excellent spatial and temporal control of light patterns, but require advanced thin film coatings to protect the implanted active devices – an ongoing research area.

Waveguide devices provide an alternative solution that allow for some of the advantages of both the simple fibre optical cannula approach and the microLED probe approach. The light source is external to the brain and is coupled via a waveguide probe to the target brain region [16, 17]. This allows for excellent spatial and temporal control of light patterns and each excitation point can illuminate a relatively large volume of tissue (∼0.1 mm^3^, suitable for primate behavioural studies). However, a significant drawback is the complicated and bulky optics required to couple light into the device [16]. The complicated optics will restrict *in vivo* experiments to either head fixed or tethered with a large fibre optic bundle.

In 2014, Pisanello *et al.* suggested a simple solution to the problem of coupling light into the brain by using tapered optical fibres [18]. This approach exploits the modal de-multiplexing properties of tapered optical fibres allowing some spatial control over the light beam. The fibre taper also allowed them to illuminate very large volumes (>10 mm^3^). Their elegant, solution still requires significant external optics and light sources that may limit behavioural studies.

As optogenetic studies progress, there is a clear need to develop new light delivery devices, specifically designed for non-human primate experiments such as those presented by Lee *et al.* [19]. A device based on the form factor of the Utah electrode array (called the Utah Optrode array - UOA) has also been developed with primate studies in mind [20]. This device allowed the coupling of light into deep brain regions with excellent spatial control. However, coupling of light into the device still required sophisticated optics such as a Spatial Light Modulator (SLM), Digital Light Projector (DLP) or the holobundle system [21]. In this paper, we propose an improvement to this device by coupling the UOA to a custom made microLED array. Coupling LED light into a multimode waveguide such as the UOA is challenging and, without complex lens systems, low coupling efficiencies (∼5%) are normal [22]. We comprehensively model the system to maximise the efficiency of coupling and develop a prototype for testing of the optical throughput and beam profiles. Thermal measurements are also made to ensure the device adheres to the strict thermal limits of an *in vivo* implant. This device has 181 individually addressable optical excitation sites capable of delivering light both superficially (81 interstitial sites) and to deep brain regions (100 needle sites). The device is capable of peak irradiances > 80 mW/mm^2^ per needle at 50 ms pulse widths with a maximum repetition rate of 1.2 Hz (to limit tissue temperature increases to less than 1 ^°^C). Furthermore, the integration of the light sources and UOA opens up the option of a fully wireless compact device [10, 15].

## 2. Methods

### 2.1 The Utah Optrode Array (UOA)

The fabricated device is a 10 x 10 array of glass needles, each measuring 1.5 mm in length and spaced at 400-micron pitch (Fig. 1 A). The needles each have a square base of side 75 μm that tapers to a sub-micron tip. A custom-designed and fabricated microLED chip is bonded directly to the backplane of the glass needle array (see Fig. 1 B). The microLED geometry can be considered as two interleaved arrays - a 10 x 10 array aligned to the glass needle shanks, and a 9 x 9 array positioned at interstitial sites between needles. The glass needles guide light to deep brain regions, while the interstitial sites couple light to surface volumes of the brain.

**Figure 1:**
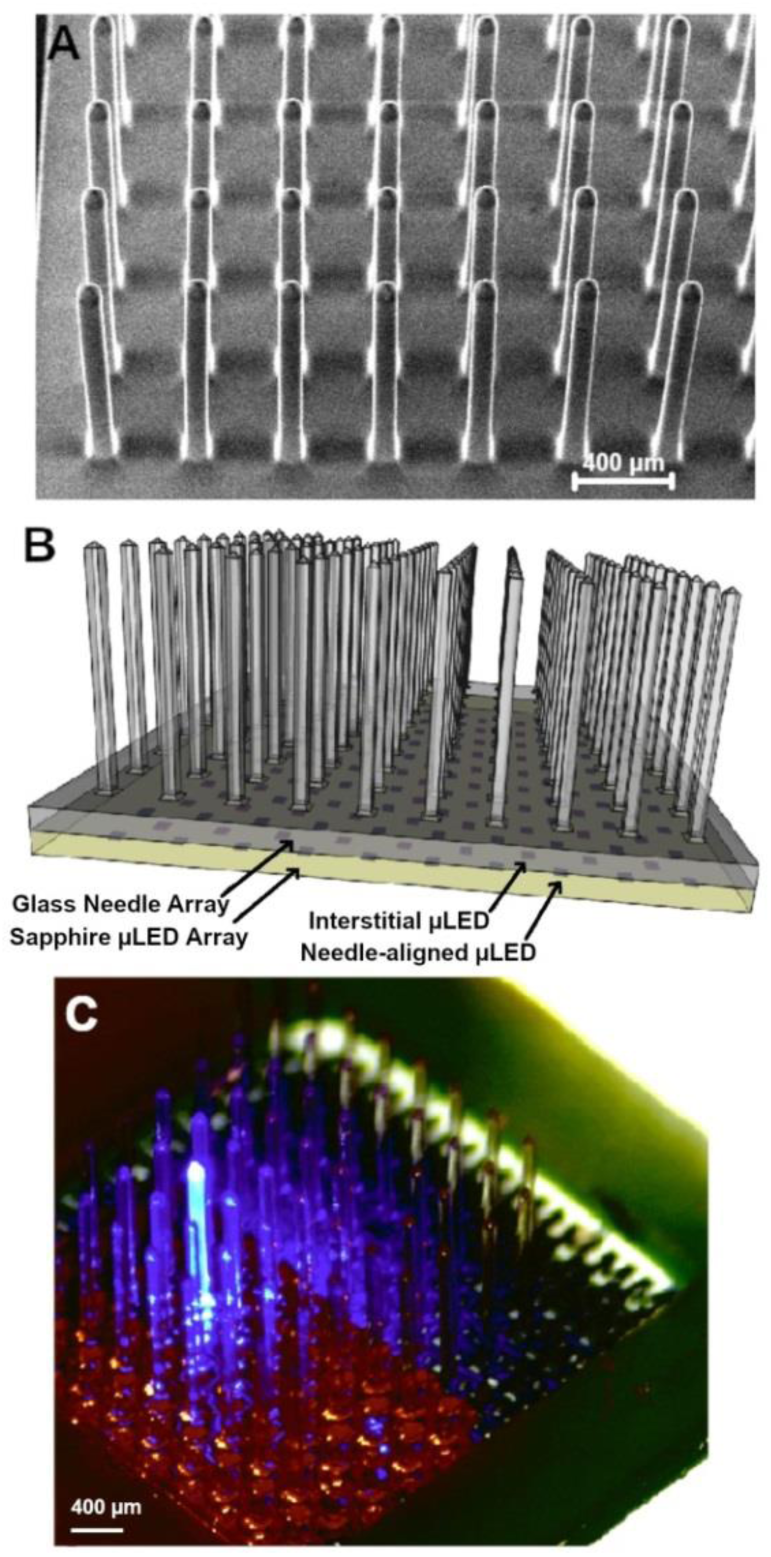
**A)** Oblique view SEM image of the Utah optrode array. **B)** Schematic of the completed device with the UOA bonded to a microLED array. The pinhole layer (not shown) is patterned onto the sapphire substrate of the microLED array before bonding. **C)** Final device including UOA with a single representative LED illuminated.

### 2.1.1 Device Microfabrication

The fabrication process flow of the UOA device is detailed in Abaya *et al*. [20]. The microLED array is fabricated from 3” GaN-on-Sapphire wafers (Xiamen Powerway Advanced Material) with a 400 μm thick sapphire substrate and a 5 μm epitaxial GaN layer stack[23]. Step 1: A thin Pd layer is deposited using electron-beam evaporation. This layer is then etched using reactive ion etching (RIE) to form the LED p-contact. Following the Pd etch, the p-GaN is etched using an inductively coupled plasma (ICP) system to create the microLED pixels (square with an 80 μm side). Step 2: The n-GaN is now patterned and etched using an ICP etch. Step 3: The sample is annealed using a rapid thermal anneal. A lift off process is then used to pattern a sputter deposited Ti/Au bilayer. Step 4: A PECVD SiO_2_ layer is deposited as an insulation layer between n-tracks and p-tracks. Vias are opened in the SiO_2_ with RIE. Step 5: The p-tracks are sputter deposited (Ti/Au) and patterned using a lift off process.

Upon completion of the microLED fabrication, the chip was mechanically thinned from the sapphire side to 150 μm and then diced into individual devices. An individual microLED array was then bonded (Norland NOA 61) to each UOA. In order to study optical cross talk between adjacent needles, a prototype device with the sapphire-side coated in a metal thin film (Ti:Au, 20nm:30nm) was fabricated. This metal layer had 40 μm diameter apertures over the microLED illumination sites – allowing light to pass through and couple into the needles while blocking stray light that would out-couple into tissue. Only half the array was covered with this pinhole layer to allow a direct comparison. Figure 1 C shows the completed device with a single needle-aligned microLED activated.

### 2.2 Optical Properties

In order to understand the optical coupling to the needle, the resultant light spread in tissue, and to optimise device design, we modelled the system using Zemax 12 in non-sequential mode. A model was developed that represented the fabricated device (Figure 2 A-B). It consisted of a GaN layer patterned into individual pixels, each with a quantum well structure where light generation occurs. To make electrical contact with the device, a thin metallic current spreading layer is placed on top of the mesa structure. Typically, this metallic layer will be semi-transparent; however, the reflectivity can be increased to close to 100% by deposition of a thicker metal layer, which is advantageous when out-coupling light via the sapphire substrate. Both semi-transparent and reflective current spreading layers are modelled.

**Figure 2:**
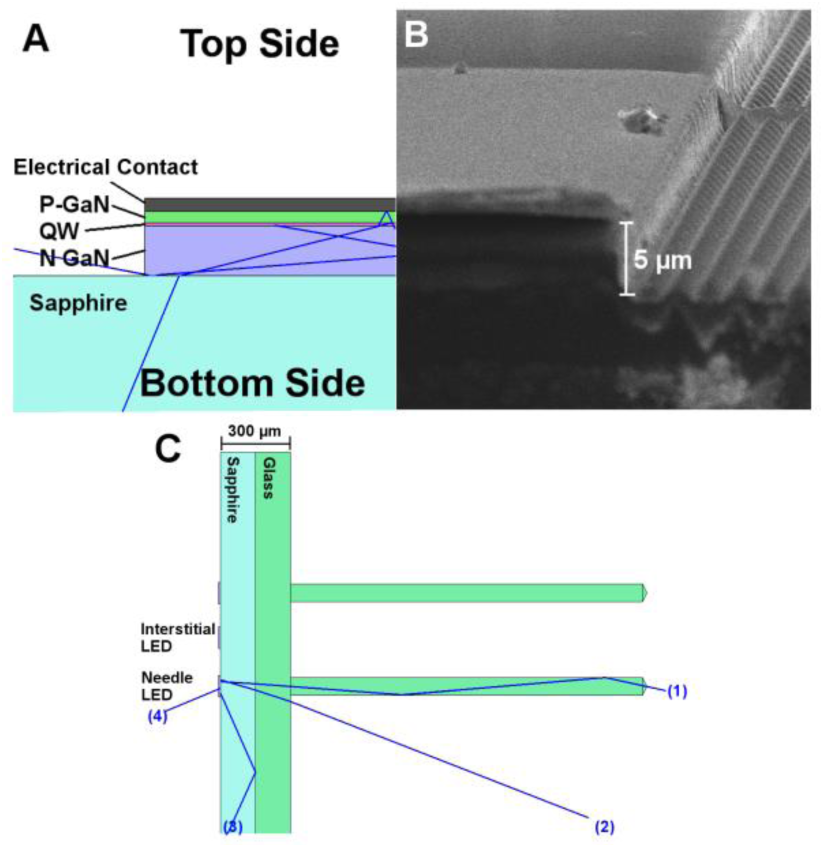
**A)** Schematic of the modelled microLED mesa structure. It consists of an electrical contact, which is designed to behave as a mirror, p-GaN layer, GaN light emitting quantum well (QW) structure, n-GaN layer and sapphire backplane. Photons are launched from a random location in the QW structure in a random direction. When they reach an interface they are reflected/refracted. **B)** SEM image of a cleaved LED mesa structure. The sapphire substrate with strain relief structure is visible on the right of the image. The GaN mesa structure is visible on the left. **C)** Schematic of the full device, light from the LED mesa was modelled propagating through the sapphire and glass backplane and into the glass needles. In this image 4 rays are shown; light coupled into the needle (1), stray light (2), trapped light (3) and back emission (4).

To model the complete device, the microLED structure, with sapphire backplane, was coupled to the needle array. The needle array can have either a glass or silicon backplane that supports the needles. The model used the peak emission wavelength of our GaN microLEDs, 450 nm. Refractive indices and absorption constants were taken from the standard Zemax libraries and the glass was modelled as BOROFLOAT^®^ 33. In each simulation run, a number of photons were launched such that > 10,000 photons crossed a modelled detector surface (thus ensuring an error of <1% from counting statistics). In subsequent analysis, an additional absorbing layer (100% absorption), with an aperture above each microLED, was also included to represent the pinhole layer mentioned in the Device Microfabrication section. An alternative to this structure is to replace the glass backplane of the needle array with a silicon substrate, which can have via structures etched in the silicon. The vias can be aligned to the microLEDs and couple light to the glass needles. This eliminates optical cross-talk, at the cost of reducing the overall light output.

The optical power of the microLEDs is measured using an optical power meter with a known collection angular aperture. Given that our devices have an electrical to light power conversion efficiency of 10-15%, typical for LEDs of this wavelength [24], we can calculate the total light generated by the QW structure for input to our models. For example, a square microLED of 80 μm side has an internal structure typically producing 80 mW optical power [25, 26].

### 2.3 Thermal Properties

Brain temperature is a carefully regulated physiological variable. Small increases in temperature (>1 ^°^C) can have profound physiological effects [27]. Since the microLED device is in physical contact with the implanted needles, thermal properties need to be understood, quantified and optimised. We measured the thermal characteristics of the complete system through thermal imaging (FLIR SC7000 series thermal camera). A calibration curve was first obtained as the temperature dependent emissivity of BOROFLOAT^®^ 33 glass is required. This was obtained by attaching a thermocouple to the device and heating to 80^°^C. Thermal images were correlated with the thermocouple temperature as it cooled from 80 ^°^C. The microLEDs were then driven at various currents, pulse widths and frequencies, and temperatures recorded from the thermal imager.

Modelling (COMSOL Multiphysics) was completed to help understand the measured thermal properties and to give an indication as to how the device will behave in brain tissue. We verified the model by comparing outputs with the experimental results in air and extended this to make predictions of temperature rises in brain tissue by altering the specific heat and thermal conductivity, taken as 3650Jkg^-1^ ^°^C^-1^ and 0.5Wm^-1^ ^°^C^-1^ for brain tissue respectively [28].

## 3. Results

A microLED yield of >95% per array was achieved, with a typical optical power – current – voltage (L-I-V) relationship shown in figure 3 A. Figure 3 B shows the peak irradiance at the tip of the needle for various currents of our prototype device. Operating at a current 100 mA the device is capable of illuminating a brain tissue volume of 0.046 mm^3^ with > 1 mW/mm^2^ irradiance (taking the measured optical output of the needle and using our optical model to determine the volume of tissue illuminated). We define a minimum threshold volume for optogenetic stimulation of 0.001 mm^3^. This is a volume of neural tissue that contains approximately 10 neurons. To illuminate this volume at > 1 mW/mm^2^ irradiance with this device requires a microLED current of 3 mA. By immersing the device in a fluorescent agarose solution it was possible to image the beam profile (Figure 3C). The measured beam width (100 μm from the tip) and decay over distance from the tip, match well with optical modelling (Figure 3C).

**Figure 3:**
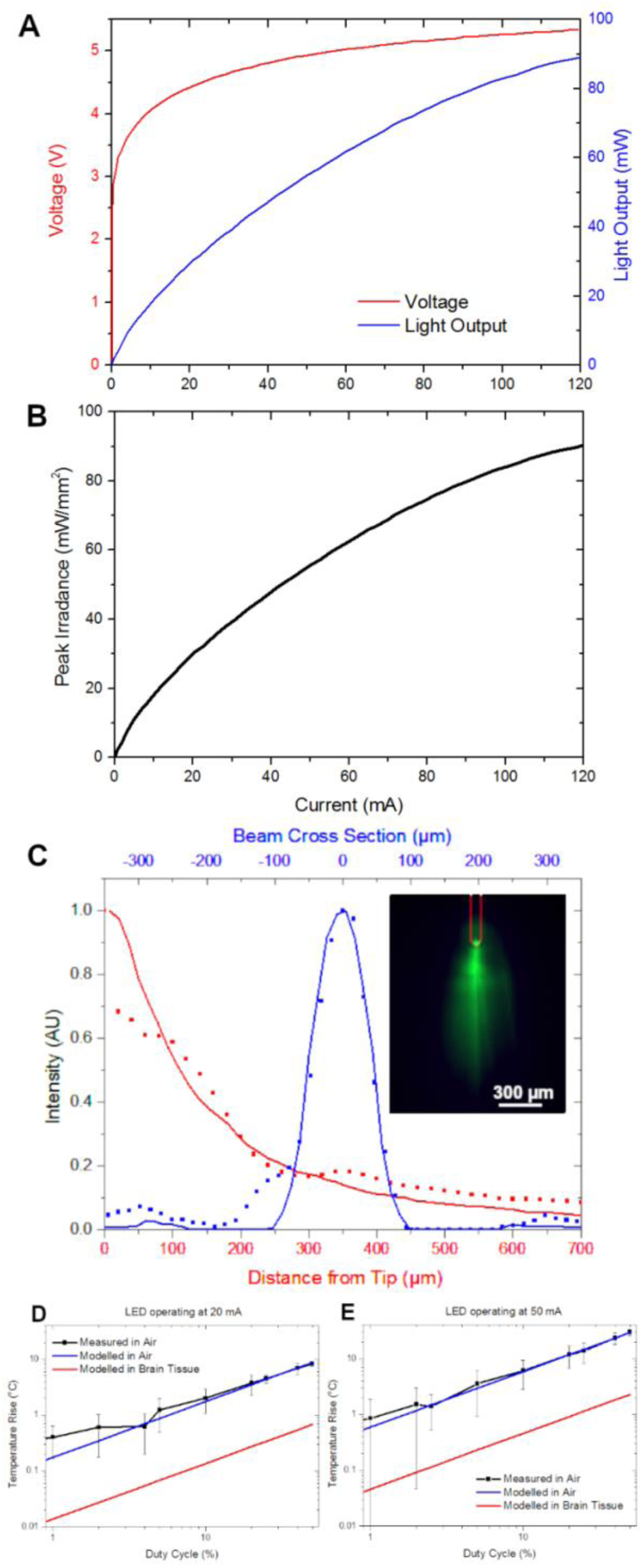
**A)** Current-Voltage and Current-Light Output characteristics of a typical microLED in our arrays. The microLEDs have a leakage current less than 80 nA (the measurement limit of our source) indicating a high quality diode. This microLED produces 80 mW of optical power at 100 mA current. **B)** The peak irradiance at the tip of the needle of our prototype device is plotted as a function of microLED current. **C)** The measured (dots) and modelled (lines) optical beam profiles from our prototype device. The beam cross section is taken 100 μm from the tip of the needle. The inset shows the beam profile as imaged in a fluorescent agarose gel. **D)** and **E)** The temperature rise for 20 mA and 50 mA LED drive current measured in air and modelled in air and brain tissue.

The device was thermally tested to assess its operating limits. Figures 3D and E show the measured thermal performance versus duty cycle of the device at 20 mA and 50 mA in air. To understand how these measurements taken in air correlate with thermal performance in brain tissue, we modelled both cases. Figures 3D and E also show a clear match between thermal modelling and measurements for air and the resultant thermal properties in brain tissue.

For integration with the Utah optrode array (UOA) device, the sapphire substrate is coupled to the glass directly. However, this compromises the efficiency of the light coupling into the needles, which is dominated by the microLED to needle-base distance. Figure 4A plots the modelled optical efficiency and power at the needle tip as the combined substrate thickness is varied. Below a substrate thickness of 40 μm the coupling efficiency is dominated by the acceptance numerical aperture of the glass needle. Our prototype device had a combined substrate thickness of 300 μm (150 μm sapphire, 150 μm glass).

**Figure 4:**
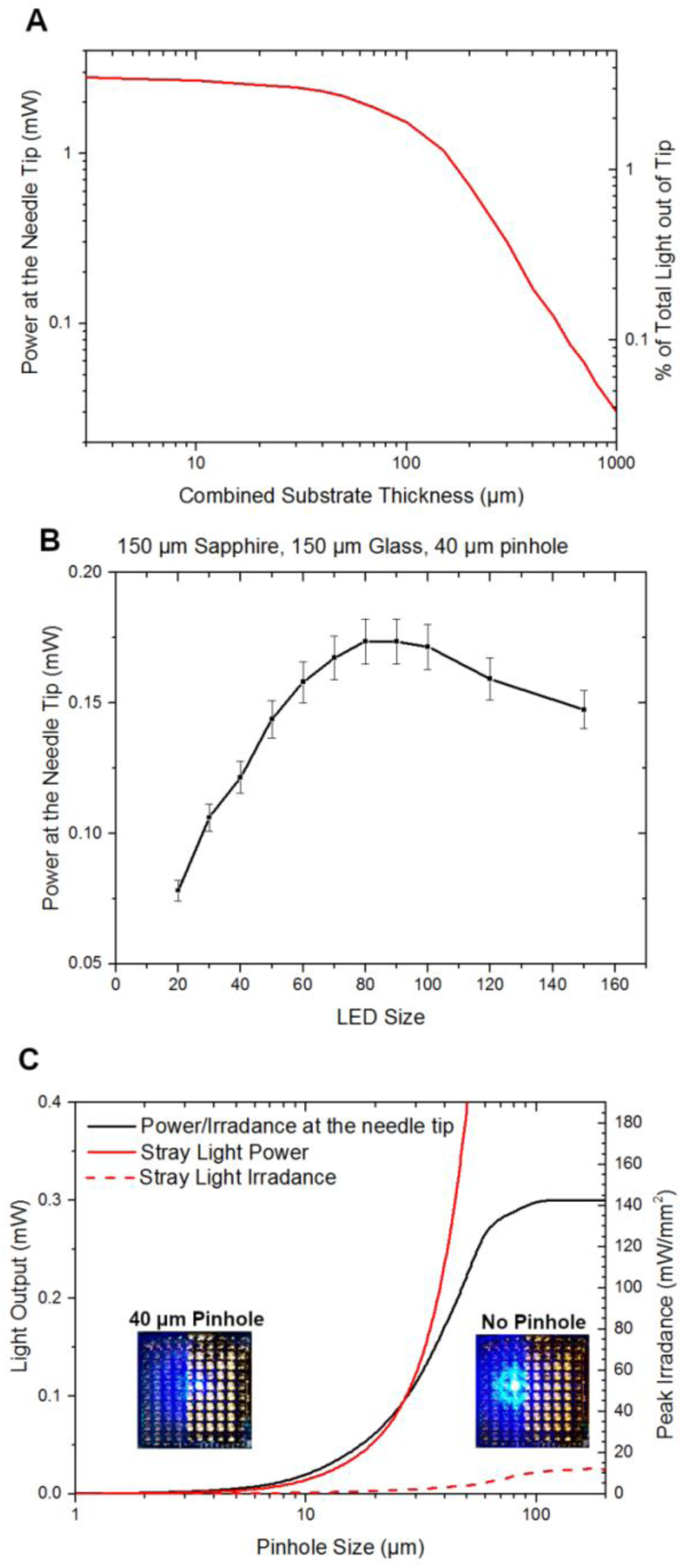
**A)** The optical power out of the tip of the needle as a function of the thickness of the substrates. The minimum total substrate thickness that we could achieve was 300 μm (150 μm sapphire, 150 μm glass). **B)** Various microLED sizes can be manufactured; the optical power at the tip of the needle for various microLED sizes is plotted. **C)** In order to minimise stray light at the base of the needles a pinhole layer is included at the interface between the microLED and UOA substrates. The power and peak irradiance at the tip of the needle (solid black line) as well as the stray light power (solid red line) and irradiance (dashed red line) are plotted as a function of pinhole size. The inserts show images of the device with a single needle aligned microLED illuminated for two cases; with a 40 μm pinhole and no pinhole.

MicroLEDs can be made in a typical size range varying from 5-200 μm [25, 26]. In figure 4B we show the optical power at the needle tip versus microLED size, with an optimum diameter from 70 – 100 μm observed.

Due to the Lambertian emission profile from LEDs [29], there will be significant amounts of stray light. This unwanted light needs to be blocked so that it does not excite optogenetic proteins away from the intended activation region. We have implemented a method to block this stray light using an absorbing layer with pinholes patterned on the interface between the microLED device and needle array. In figure 4C, square optical pinholes of various dimensions ranging from 1 μm to 200 μm were modelled. Photons were detected on two modelled detector surfaces, 100 μm from the base of the needle and at the tip of the needle. For pinhole sizes less than 10 μm, light was only coupled into the needle. However, this size limits the light output to less than 10 mW/mm^2^. As the pinhole size was increased, more stray light coupled out around the base of the needle. To allow for sufficient light at the tip of the needle a pinhole size of 40 μm was deemed optimal. The measured and modelled optical properties of our prototype device are summarised in table 1. To put these figures in perspective, to achieve a comparable illumination volume to microLED probe devices (0.01 mm^3^) [12-14] a coupling efficiency of 0.05% would be required. To achieve an illumination volume of > 0.1 mm^2^ (similar to typical fibre optic cannula) a coupling efficiency of > 0.5% would be required.

**Table 1:** Modelled and measured performance of the integrated device. Here, light emission is through the sapphire substrate of the microLED array and passes through the backplane of the UOA before coupling into the glass needles. A metal plane with pinhole apertures is also incorporated between the sapphire and glass.

| Distance between microLED and needle base | Modelled Light Out of tip | Modelled Stray Light | Trapped or Absorbed Light | Measured Light Output @ 100 mA |
| --- | --- | --- | --- | --- |
| 300 $\mu\text{m}$ (no pinhole) | 0.4 % (330$\mu\text{W}$) | 16.1 % | 79.3 % | 0.45% (360 $\mu\text{W}$) |
| 300 $\mu\text{m}$ (with 40$\mu\text{m}$ pinhole) | 0.2 % (170$\mu\text{W}$) | 0.3 % | 95.3 % | 0.22% (179 $\mu\text{W}$) |

The described optical model agrees well with experiment and illustrates the problems of efficiently coupling light into the needles, which is a classic problem with incoherent light sources [22]. Next, we have used the model to optimise the optical efficiency of the system, within the fabrication constraints of our device. During microLED fabrication, a Pd current spreading layer is deposited to make electrical contact with the microLED. Ordinarily this layer is thin enough to be semi-transparent. To increase the optical efficiency of the microLED device an additional Ti/Au layer is added to the model to act as a mirror layer and reduce emission from the device in an undesired direction, figure 5 A. However, the optical efficiency of this optogenetic device is mostly determined by the backplane (sapphire and glass) thickness, so top-emitting (through the p-contact) microLEDs are also considered (figure 5 B). This would allow significant reduction in the backplane thickness of the device.

**Figure 5:**
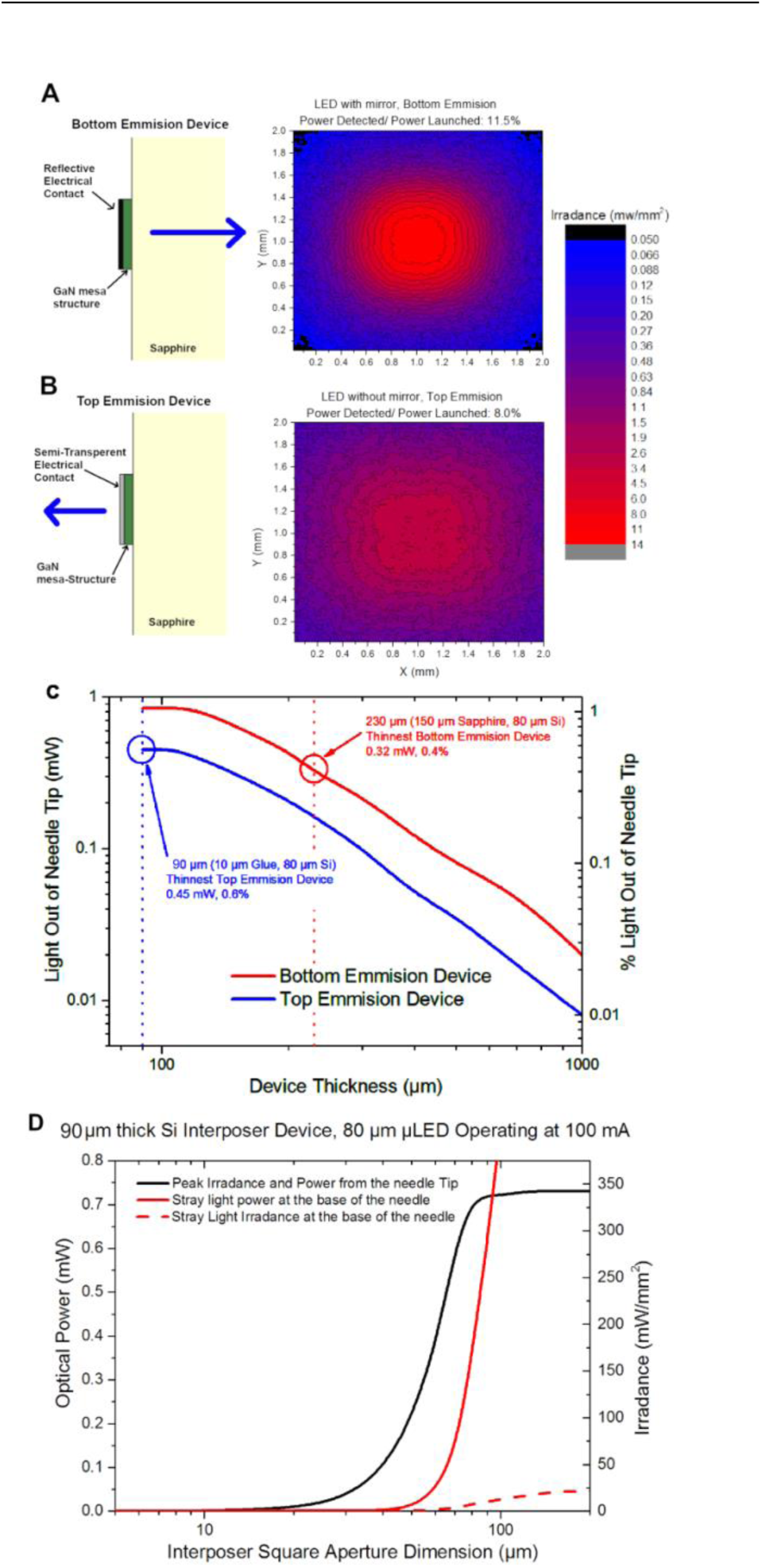
**A)** A typical GaN on Sapphire microLED is fabricated so that the electrical contact behaves as a mirror to maximise light output through the sapphire. The modelled beam profile from an 80 μm bottom emission microLED is shown. **B)** It is also possible to produce a top emitting microLED using transparent electrical contacts. The modelled beam profile from an 80 μm top emission microLED is shown. **C)** It is possible to fabricate the UOA on a Si substrate (as opposed to glass) with optical interposers etched to allow light to pass through. We model the optical power out of the tip of a needle for both top and bottom emission microLEDs as a function of substrate thickness. Highlighted are the limits of our current fabrication technology. **D)** The power and peak irradiance at the tip of the needle (solid black line) as well as the stray light power (solid red line) and irradiance (dashed red line) are plotted as a function of Si interposer dimension.

Recent developments in the production of the UOA allow for the glass backplane to be replaced with Silicon. Using conventional micro-processing techniques, holes can be opened in the Si backplane [30], creating an optical interposer. This Si backplane can also be mechanically thinned to approximately 80 μm. The device backplane thickness versus optical power of our silicon backplane device is modelled in figure 5 C for both a top and bottom emitting microLED device. Interposer aperture size was also modelled to determine the optimal dimension. Figure 5 D shows the effect of increasing the interposer aperture dimension on both the light out of the tip and stray light (for a top emission device with 10 μm optical adhesive and 80 μm thick Si interposer layer). An interposer aperture size of 60 μm completely blocks stray light while allowing 0.38 mW out of the tip of the needle, this corresponds to a peak irradiance of 184 mW/mm^2^.

A full summary of the device optimisations is shown in table 2. We have identified two clear opportunities to improve the optical efficiency of the device: 1) a bottom emission device (with 150 μm sapphire backplane) directly integrated with an UOA that has an 80 μm thick Si backplane. This device will give an optical efficiency of 0.4% with no stray light. 2) A top emission LED device, bonded to an 80 μm thick Si backplane needle array and under-filled with an optical adhesive (giving a total backplane thickness of 90 μm). This device would give an optical efficiency of 0.6% with negligible stray light.

**Table 2:** Summary of the modelled optical efficiencies of the proposed devices. Highlighted in grey, are the two devices that have been fabricated. For reference, if 1% of the light was coupled to the needle, it would correspond to a peak irradiance in the tissue of ∼ 300 mW/mm^2^. In the bottom emission case 4.2 % of the light is emitted in the opposite direction of the needle. In the top emission cases 9.5 % of the light is emitted in the opposite direction of the needle.

| Device description | Light from needle (%) | Stray light (%) | Trapped or absorbed light (%) |
| --- | --- | --- | --- |
| Bottom emission, 150$\mu\text{m}$ sapphire, 80$\mu\text{m}$ silicon, 60$\mu\text{m}$ interposer hole | 0.4 % | 0 % | 95.4 % |
| Top emission, 10$\mu\text{m}$ optical adhesive, 150$\mu\text{m}$ glass, no pinhole | 0.3 % | 15.6 % | 74.6 % |
| Top emission, 10$\mu\text{m}$ optical adhesive, 150$\mu\text{m}$ glass, 40$\mu\text{m}$ pinhole | 0.2 % | 1.5 % | 88.8% |
| Top emission, 10$\mu\text{m}$ optical adhesive, 80$\mu\text{m}$ silicon, 60$\mu\text{m}$ interposer hole | 0.6 % | 0.1 % | 89.8 % |

## 4. Discussion

There is a clear need for an optogenetic device that can deliver structured light to deep brain structures in larger mammals. The novel microLED-UOA neurophotonic technology introduced here is capable of providing sufficient light to excite hundreds of neurons per needle site in non-human primates or other large mammals. The integrated nature of this electrically addressable device has the potential to enable wireless operation and so open the technology to wider neuroscience applications. In parallel with this, the fabrication process is highly scalable and should now allow for the rapid production and distribution of the devices.

Other options for coupling light into UOA device have been considered but these have been ruled out for various reasons. A free space optics microLED array based system could give very high coupling efficiency (>2.5%) but optical alignment would be extremely challenging, ruling out chronic and freely behavioural studies. Other optical sources have also been considered. VCSEL arrays [31] have recently become available, however, these devices do not have the fabrication maturity of microLED arrays and, although they will give a device with greater optical efficiency, this comes at the cost of a less power efficient light source (worse thermal performance) at the low optical powers required.

Comprehensive electrical, optical and thermal testing of the microLED-UOA device was performed, alongside extensive modelling. Optical cross-talk between the glass needles can reach significant levels if strategies to block stray light are not adopted. The simplest of such was a metallic absorbing layer with small apertures, aligned with the microLEDs, and deposited on the sapphire side of the microLED chip. The final prototype device could emit 170 μW of light at 450 nm wavelength from the tip of each needle, when driven at 100 mA. This gave a peak irradiance of 81 mW/mm^2^ at the tip of the needle, modelling indicates that this is sufficient to optogenetically excite a volume of tissue of 0.046 mm^3^, containing approximately 5500 neurons (approximating neuronal density to 1.2 x 10^5^ neurons/mm^3^ [32]). Thermal measurements and modelling allow us to place a duty cycle limit on the operation of the device to ensure that neural tissue does not exceed a 1 ^°^C temperature rise. This limit is 75 % for a microLED operating at 20 mA, 22% at 50 mA and 6% at 100 mA.

Modelling also allowed the optical design to be optimised. The pinhole structure used to block stray light in our prototype device is not ideal. It cannot block all stray light and still allow for enough light at the needle tip. An alternative option would be to replace the glass backplane of the needle array with a silicon backplane. Optical interposers can be opened in this backplane allowing light to be coupled into needle and interstitial sites. This silicon-interposer design has the added advantage that it can be thinned to 80 μm while still allowing for a stable device with high yield. Thinning of the device would allow for the biggest gain in efficiency. However, thinning of the sapphire backplane of the microLED device to below 150 μm significantly reduces device yield. By switching to a top emitting LED we can design a device that has an effective backplane of 10 μm (optical adhesive) at the expense of reduced microLED light output. Combining these two approaches we could potentially increase the optical efficiency of our device by 3-fold (from 0.2% to 0.6%), corresponding to a peak irradiance of 184 mW/mm^2^ and increasing the maximum illumination volume to >0.1mm^3^, while also ensuring there is no optical cross talk. Furthermore, by increasing the optical efficiency we will be able to significantly reduce the power to each LED and hence reduce the operating temperature. An increase in optical efficiency from 0.2 % to 0.6 % would reduce the power required for each microLED from 0.5 W to 0.113 W for a peak irradiance of > 80 mW/mm^2^. This would mean that operating with a 50 ms pulse width the maximum repetition rate would increase from 1.2 Hz to 10 Hz.

## Acknowledgements

This work was supported by the NIH BRAIN Initiative Program, through grant U01 NS099702.

## References

1. Deisseroth, K., Optogenetics: 10 years of microbial opsins in neuroscience. Nature Neuroscience, 2015. 18: p. 1213.

2. Hunnicutt, B.J., et al., A comprehensive thalamocortical projection map at the mesoscopic level. Nature Neuroscience, 2014. 17: p. 1276.

3. Kress, G.J., et al., Convergent cortical innervation of striatal projection neurons. Nature Neuroscience, 2013. 16: p. 665.

4. Domingos, A.I., et al., Leptin regulates the reward value of nutrient. Nature Neuroscience, 2011. 14: p. 1562.

5. Han, X., Optogenetics in the nonhuman primate. Progress in brain research, 2012. 196: p. 215–233.

6. Prakash, R., et al., Two-photon optogenetic toolbox for fast inhibition, excitation and bistable modulation. Nature Methods, 2012. 9: p. 1171.

7. Hososhima, S., et al., Near-infrared (NIR) up-conversion optogenetics. Scientific Reports, 2015. 5: p. 16533.

8. Chernov, K.G., et al., Near-Infrared Fluorescent Proteins, Biosensors, and Optogenetic Tools Engineered from Phytochromes. Chemical Reviews, 2017. 117(9): p. 6423–6446.

9. Zhang, F., et al., Optogenetic interrogation of neural circuits: technology for probing mammalian brain structures. Nature Protocols, 2010. 5: p. 439.

10. Gagnon-Turcotte, G., et al., A Wireless Optogenetic Headstage with Multichannel Electrophysiological Recording Capability. Sensors (Basel, Switzerland), 2015. 15(9): p. 22776–22797.

11. Buzsáki, G., et al., Tools for probing local circuits: high-density silicon probes combined with optogenetics. Neuron, 2015. 86(1): p. 92–105.

12. McAlinden, N., et al., Optogenetic activation of neocortical neurons in vivo with a sapphire-based micro-scale LED probe. Frontiers in Neural Circuits, 2015. 9(25).

13. Scharf, R., et al., Depth-specific optogenetic control in vivo with a scalable, high-density µLED neural probe. Scientific Reports, 2016. 6: p. 28381.

14. Wu, F., et al., Monolithically Integrated μLEDs on Silicon Neural Probes for High-Resolution Optogenetic Studies in Behaving Animals. Neuron, 2015. 88(6): p. 1136–1148.

15. Kim, T.-i., et al., Injectable, Cellular-Scale Optoelectronics with Applications for Wireless Optogenetics. Science, 2013. 340(6129): p. 211.

16. Zorzos, A.N., E.S. Boyden, and C.G. Fonstad, Multiwaveguide implantable probe for light delivery to sets of distributed brain targets. Optics Letters, 2010. 35(24): p. 4133–4135.

17. Hoffman, L., et al. High-density optrode-electrode neural probe using SixNy photonics for in vivo optogenetics. in 2015 IEEE International Electron Devices Meeting (IEDM). 2015.

18. Pisanello, F., et al., Multipoint-Emitting Optical Fibers for Spatially Addressable In Vivo Optogenetics. Neuron, 2014. 82(6): p. 1245–1254.

19. Lee, J., et al., Transparent intracortical microprobe array for simultaneous spatiotemporal optical stimulation and multichannel electrical recording. Nature Methods, 2015. 12: p. 1157.

20. Abaya, T.V.F., et al., A 3D glass optrode array for optical neural stimulation. Biomedical Optics Express, 2012. 3(12): p. 3087–3104.

21. Boutte, R.W., et al. Utah optrode array customization using stereotactic brain atlases and 3-D CAD modeling for optogenetic neocortical interrogation in small rodents and nonhuman primates. 2017. SPIE.

22. Keiser, G., Optical Fiber Communications. 4th Ed ed. 2008, New York: McGraw-Hill Education.

23. Ryou, J.-H. and W. Lee, GaN on sapphire substrates for visible light-emitting diodes, in Nitride Semiconductor Light-Emitting Diodes (LEDs) (Second Edition), J. Huang, H.-C. Kuo, and S.-C. Shen, Editors. 2018, Woodhead Publishing. p. 43–78.

24. Greshnov, A.A., et al., Comparative study of quantum efficiency of blue LED with different nanostructural arrangement. physica status solidi c, 2007. 4(8): p. 2981–2985.

25. Tian, P., et al., Size-dependent efficiency and efficiency droop of blue InGaN micro-light emitting diodes. Applied Physics Letters, 2012. 101(23): p. 231110.

26. McKendry, J.J.D., et al., Visible-Light Communications Using a CMOS-Controlled Micro-Light-Emitting-Diode Array. Journal of Lightwave Technology, 2012. 30(1): p. 61–67.

27. Andersen, P. and E.I. Moser, Brain temperature and hippocampal function. Hippocampus, 1995. 5(6): p. 491–498.

28. Maged, M.E., et al., Bio-heat transfer model of deep brain stimulation-induced temperature changes. Journal of Neural Engineering, 2006. 3(4): p. 306.

29. Griffin, C., et al., Beam divergence measurements of InGaN/GaN micro-array light-emitting diodes using *confocal microscopy*. Applied Physics Letters, 2005. 86(4): p. 041111.

30. Scharf, R., et al. A compact integrated device for spatially-selective optogenetic neural stimulation based on the Utah Optrode Array. in SPIE BiOS. 2018. SPIE.

31. Mei, Y., et al., Quantum dot vertical-cavity surface-emitting lasers covering the ’green gap’. Light: Science & Applications, 2017. 6: p. e16199.

32. Collins, C.E., et al., Neuron densities vary across and within cortical areas in primates. Proceedings of the National Academy of Sciences, 2010. 107(36): p. 15927–15932.

